# Type I and III IFNs produced by the nasal epithelia and dimmed inflammation are key features of alpacas resolving MERS-CoV infection

**DOI:** 10.1101/2020.12.22.423939

**Authors:** Nigeer Te, Jordi Rodon, Maria Ballester, Mónica Pérez, Lola Pailler-García, Joaquim Segalés, Júlia Vergara-Alert, Albert Bensaid

## Abstract

While MERS-CoV (Middle East respiratory syndrome Coronavirus) provokes a lethal disease in humans, camelids, the main virus reservoir, are asymptomatic carriers, suggesting a crucial role for innate immune responses in controlling the infection. Experimentally infected camelids clear infectious virus within one week and mount an effective adaptive immune response. Here, transcription of immune response genes was monitored in the respiratory tract of MERS-CoV infected alpacas. Concomitant to the peak of infection, occurring at 2 days post inoculation (dpi), type I and III interferons (IFNs) were maximally transcribed only in the nasal mucosa of alpacas, provoking the induction of interferon stimulated genes (ISGs) along the whole respiratory tract. Simultaneous to mild focal infiltration of leukocytes in nasal mucosa and submucosa, upregulation of the anti-inflammatory cytokine IL10 and dampened transcription of pro-inflammatory genes under NF-κB control were observed. In the lung, early (1 dpi) transcription of chemokines (CCL2 and CCL3) correlated with a transient accumulation of mainly mononuclear leukocytes. A tight regulation of IFNs in lungs with expression of ISGs and controlled inflammatory responses, might contribute to virus clearance without causing tissue damage. Thus, the nasal mucosa, the main target of MERS-CoV in camelids, is central in driving an efficient innate immune response based on triggering ISGs as well as the dual anti-inflammatory effects of type III IFNs and IL10.

**Author summary:** Middle East respiratory syndrome coronavirus (MERS-CoV) is the etiological agent of a respiratory disease causing high mortality in humans. In camelids, the main MERS-CoV reservoir host, viral infection leads to subclinical disease. Our study describes transcriptional regulations of innate immunological pathways underlying asymptomatic clinical manifestations of alpacas in response to MERS-CoV. Concomitant to the peak of infection, these animals elicited a strong transient interferon response and induction of the anti-inflammatory cytokine IL10 in the nasal mucosa. This was associated to a dimmed regulation of pro-inflammatory cytokines and induction of interferon stimulated genes along the whole respiratory mucosa, leading to the rapid clearance of the virus. Thus, innate immune responses occurring in the nasal mucosa appear to be the key in controlling MERS-CoV disease by avoiding a cytokine storm. Understanding on how asymptomatic host reservoirs counteract MERS-CoV infection will aid in the development of antiviral drugs and vaccines.

## Introduction

The Middle East respiratory syndrome (MERS) is a disease caused by a zoonotic Coronavirus (MERS-CoV) that emerged in 2012 in the Kingdom of Saudi Arabia [1] raising a toll of 2,562 confirmed human cases in 27 countries, with 881 deaths until the November 2020 [2]. In humans, MERS-CoV infection ranges from asymptomatic to severe or even fatal respiratory disease [3]. Dromedary camels are the main viral reservoir [4], and all camelids are susceptible to the virus, under both natural and experimental conditions [5–10]. However, despite consequent tissue viral loads and high viral shedding at the upper respiratory tract (URT) level, infection in camelids is asymptomatic, leading to a rapid clearance of the virus [5,9,11] and the establishment of a solid acquired immunity. Indeed, field studies revealed a high proportion of serum neutralizing antibodies in dromedary camels [8,12]. Innate immune responses are essential as they link adaptive immunity [13] and are key players in the pathology of diseases [14]. Nevertheless, the severity of MERS lesions in humans has been attributed to aberrant innate and adaptive immune responses based essentially on data obtained from macrophages isolated from healthy donors or infected patients, as well as dosage of cytokines/chemokines from bronchoalveolar lavages. The outcome of these studies reveals an overproduction of proinflammatory cytokines/chemokines due to the activation of C-type lectin receptors, RIG-I like receptors (RLRs) and an impaired production of type I interferons (IFN) [15–18]. High production and persistent high levels of these cytokines in macrophages are likely to exacerbate disease severity. Despite that macrophages can be infected, at least *in vitro,* MERS-CoV has a preferable tropism for respiratory epithelial cells [18]. To date, no data is available on innate immune responses affecting the human respiratory mucosa *in vivo* and most information was mainly obtained from *ex vivo* pseudostratified primary bronchial airway epithelial cells [18,19], immortalized epithelial cells [19,20] or respiratory explants [21–23]. Although some contradictory results have been reported, MERS-CoV infections in these cells or tissues led to the conclusion that type I and III IFN are inhibited [24,25] or, when delayed [26], weakly expressed [27]. Nonetheless, some MERS-CoV African strains isolated from dromedary camels, as opposed to the Arabian human isolated EMC/2012 strain, can induce higher levels of IFNß, IFNλ1, IFN stimulated genes (ISGs) and proinflammatory cytokines mRNA in Calu-3 cells after 48 h of infection [22]. Indeed, the prototypic strain EMC/2012 and MERS-CoV African strains belong to different clades (A and C respectively) displaying deletions particularly in the ORF4b [22]. In that respect, several viral factors, including MERS-CoV accessory proteins 4a and 4b, have been shown in respiratory epithelial cell lines to antagonize the production of IFNs, to interfere with the NF-κB signaling pathway by avoiding production of proinflammatory cytokines [28] and inhibiting the protein kinase R-mediated stress response [29]. In addition, in epithelial cell lines, the structural viral protein N interacts with the host E3 ligase tripartite motif protein 25 (TRIM25) impeding ubiquitination of RIG-I and further expression of type I interferons and IFN-λ1 [30]. Therefore, in contrast to the cytokine storm provoked by macrophages, epithelial innate immune responses seem to be profoundly paralyzed. This is of major consequence for the progression of the disease, since the respiratory mucosa is the primary barrier for MERS-CoV. Recent findings indicate that type III IFNs confer initial protection, restricting tissue damage at the mucosa level by limiting inflammatory responses and potentiating adaptive immunity. Furthermore, it is postulated that when the viral burden is high and the mucosal fitness is broken, type I IFNs take over, leading to enhanced immune responses provoking also uncontrolled pro-inflammatory responses [31].

Owing that bats, the primary reservoir for coronaviruses, are tolerant to MERS-CoV (and other viruses) due to a dampened Nod-like receptor family pyrin domain-containing 3 (NLRP3) inflammasome [32], it was of major interest to gain insights into innate immune responses induced by MERS-CoV in camelids. Therefore, by understanding how reservoir/intermediary hosts control MERS-CoV and by extension other coronaviruses, a wealth of information could be translated to other species experimenting severe disease for the improvement of prevention, treatments and vaccines.

In the present study, alpacas were experimentally infected and monitored at the transcriptional level for a set of innate immune response genes along the upper and lower respiratory tract (LRT) during four consecutive days. Special attention was given to the nasal epithelium as it is the primary site of MERS-CoV replication. Regulation of IFNs, pattern recognition receptors (PRRs), IFN regulatory transcription factors (IRFs) and enzymes constituting the NLRP3 inflammasome or the NF-κB pathway was analyzed providing insights on signaling mechanisms important for MERS-CoV replication and disease progression, elucidating key regulatory host factors to counteract MERS-CoV infection.

## Results

### Clinical signs of alpacas following MERS-CoV infection

The objective of this study was to evaluate early innate immune responses upon MERS-CoV infection in camelids. The MERS-CoV Qatar15/2015 strain was selected because clade B strains are nowadays exclusively circulating in the Arabian Peninsula [4,33]. Three alpacas (AP13-AP15) were euthanized before virus inoculation and served as controls, while groups of three animals (AP1-AP3, AP4-AP6, AP7-AP9, AP10-AP12) were sacrificed sequentially during four consecutive days after MERS-CoV inoculation. nasal swabs (NS) and respiratory tissue samples were collected at the day of euthanasia.

Following virus inoculation, only one alpaca (AP6) secreted a mild amount of nasal mucus on 2 days post inoculation (dpi). None of the remaining animals showed clinical signs and basal body temperatures remained constant (below 39.5°C) throughout the study.

### Nasal viral shedding and MERS-CoV loads in respiratory tracts during infection in alpacas

On the day of euthanasia (0, 1, 2, 3, and 4 dpi), NS were collected. Viral detection and loads were assessed by RT-qPCR. All MERS-CoV inoculated alpacas (AP1-AP12) shed viral RNA, but no major differences were detected between the different time points. Also, viral titration in NS showed that all inoculated animals excreted infectious MERS-CoV. Maximal viral loads in the nasal cavity were reached at 2 dpi. Of note, animal AP6 shed the highest loads of infectious virus (4.8 TCID50/ml). None of the alpacas, including negative controls (AP13-AP15), had viral RNA or infectious virus on 0 dpi (Fig 1A). MERS-CoV RNA was detected in all homogenized respiratory tissues during infection. The higher viral loads were found on 2 dpi in nasal turbinates and bronchus with a significant increase compared to those detected on 1 dpi. Trachea and lung displayed the lowest viral RNA loads. In all cases, at 4 dpi, infectious virus was still excreted, and viral RNA was present in the respiratory tract (Fig 1B).

**Fig 1.**
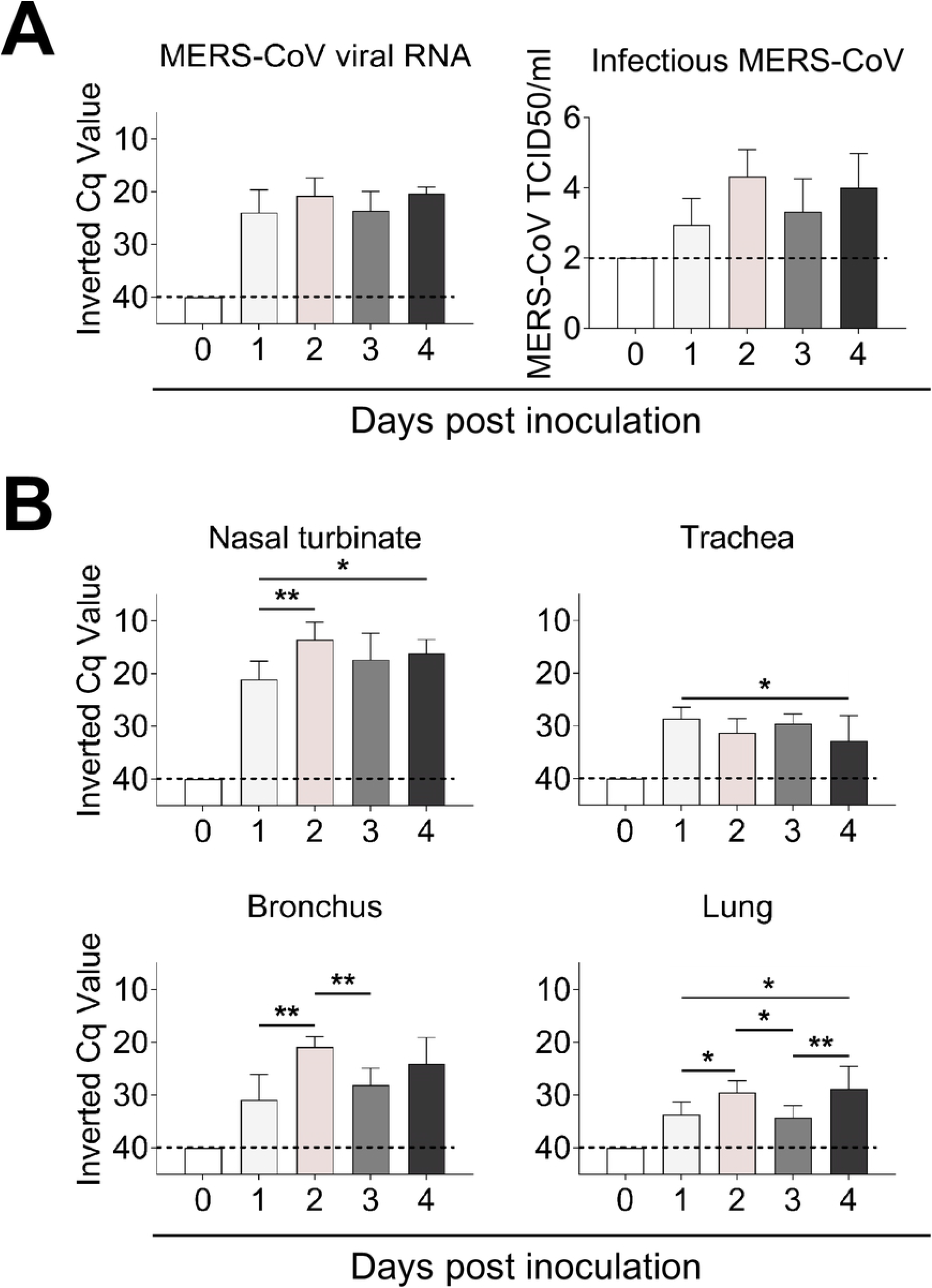
Viral loads in nasal swabs and respiratory tissues of MERS-CoV-infected alpacas. (A) Viral RNA (left) and infectious MERS-CoV (right) loads from nasal swab samples collected at the day of euthanasia. (B) Viral RNA from respiratory tissues of alpacas collected at different dpi. Viral loads were determined by the UpE real-time RT-qPCR (A and B). Each bar represents the mean Cq value +SD of infected tissues from 3 animals euthanized on 0, 1, 2, 3 and 4 dpi, respectively (in total 15 animals). Statistical significance was determined by Tukey’s multiple comparisons test. \**P* < 0.05; \*\**P* < 0.01; \*\*\**P* < 0.001. Dashed lines depict the detection limit of the assays. Cq, quantification cycle; TCID50, 50% tissue culture infective dose.

### MERS-CoV establishes early infections in URTs and LRTs of alpacas

Histopathology of URTs and LRTs was assessed in a blinded manner independently by veterinary pathologists. On 2 and 3 dpi, histological lesions in MERS-CoV infected alpacas were limited to the respiratory tract, being of multifocal distribution and mild. Lesions in nasal turbinates were characterized by mild rhinitis, segmental hyperplasia of the nasal epithelium, lymphocytic exocytosis, loss of epithelial polarity and tight junction integrity. Additionally, small numbers of lymphocytes with fewer macrophages infiltrated the underlying submucosa. No microscopic lesions were observed in nasal turbinates on 0, 1 and 4 dpi in any of the animals, which displayed a multifocal localization of the MERS-CoV antigen detected by immunohistochemistry (IHC). While only a few pseudostratified columnar epithelial cells in the nose contained MERS-CoV antigen on 1 dpi, the number of positive epithelial cells was remarkably high on 2 dpi; such number steadily decreased in the alpacas necropsied on the following days. On 4 dpi, the MERS-CoV antigen was scarcely detected with no evidence of microscopic lesions (Fig 2A).

**Fig 2.**
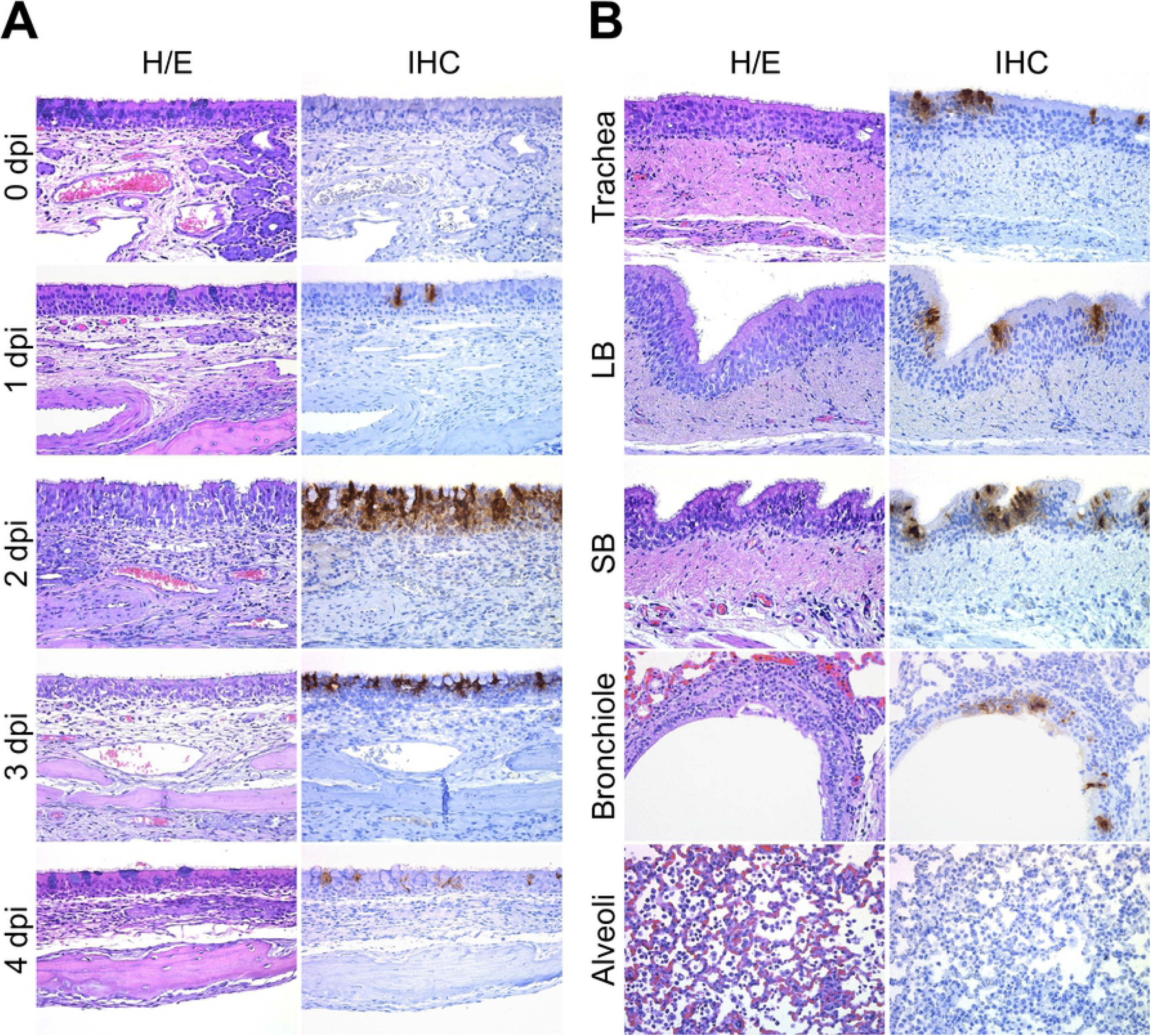
Histopathological changes and viral detection in respiratory tracts of alpacas inoculated with MERS-CoV. All respiratory tissues were fixed in 10% neutral-buffered formalin. (A) Nasal turbinate tissue sections of alpacas from the non-infected group (0 dpi), and from those necropsied on1 to 4 dpi. (B) Tissue sections of trachea, bronchus and lung (bronchiole and alveoli) from infected alpacas at 2 dpi. Original magnification: ×400 for all tissues. See S1 Table for the detailed distribution of MERS-CoV antigen in respiratory tracts. Abbreviations: H/E, hematoxylin and eosin stain; IHC, immunohistochemistry; LB, large bronchus; SB, small bronchus.

Trachea and bronchus showed multifocal, mild tracheitis/bronchitis, with the presence of few lymphocytes in the epithelium and mild infiltration of the submucosa by lymphocytes and macrophages on 2 dpi. In line with these observations, MERS-CoV infected cells were rarely detected within the epithelium on 2 dpi by IHC, mostly in areas displaying these minimal lesions (Fig 2B). Remarkably, the trachea of AP6 harbored the highest viral load as detected by IHC. Few labeled cells were observed on 4 dpi (S1 Table) with no evidence of lesions. Additionally, lung lobes showed mild multifocal perivascular and peribronchiolar infiltration by lymphocytes and the presence of monocyte/macrophages within the alveoli, being more evident on 2dpi. MERS-CoV antigen was occasionally observed in bronchiolar epithelial cells on 2 dpi (Fig 2B), and rarely on 4 dpi (S1 Table). Of note, pneumocytes were not labeled by MERS-CoV IHC (Fig 2B).

### Viral loads in micro-dissected nasal tissues and whole tracheal and lung sections

Viral loads and viral transcription/replication were also assessed on methacarn-fixed paraffin-embedded (MFPE) tissue sections which were further used for cytokine quantification. Special attention was given to the nasal epithelium as it is the privileged tissue for viral replication. Thus, laser capture microdissection (LCM) was used to obtain, when possible and for each animal, nasal epithelial areas positive or negative for MERS-CoV, as assessed by IHC, in different but consecutive (parallel) MFPE tissue sections. Their respective underlying submucosa were also collected at 1 and 3 dpi (S1 Fig). Moreover, MFPE sections of tracheas and lungs were directly scrapped from the slide. On 1 dpi only a few isolated nasal epithelial cells were IHC positive. These were microdissected, as described above, with the surrounding IHC negative cells to get enough RNA for the microfluidic PCR quantitative assay. To the contrary, due to the massive infection in nasal tissues at 2 dpi, only IHC negative nasal epithelial areas from AP4 could be collected. Viral loads in micro-dissected nasal tissues and MFPE tracheal and lung scrapped samples were quantified with UpE primers using the microfluidic quantitative PCR assay. As illustrated in A in S2 Fig, micro-dissected MERS-CoV infected epithelial areas, as assessed by their parallel IHC stained sections, showed higher viral loads than non-labelled epithelial areas, confirming the validity of the technique. In agreement with immunohistochemical observations, MERS-CoV RNA was of lower abundance in submucosal layers. As expected, and according to results obtained with tissue macerates (Fig 1B), microfluidic quantification of MERS-CoV viral RNA revealed a much lower degree of virus replication in trachea and lung than in nasal tissues. These results were confirmed with the M mRNA microfluidic PCR (B in S2 Fig and A in S3 Table) and, as expected, showed a greater extension of MERS-CoV infection in tissues, in particular in the nasal epithelia, than that revealed by IHC.

### High induction of type I and III IFNs in the nasal mucosa occurs at the peak of infection in alpacas

In order to investigate the antiviral pathways induced upon MERS-CoV infection, relative mRNA expression levels for 37 innate immune response genes were assessed along the respiratory tract with the same cDNA samples as those used for viral UpE and M mRNA quantifications in a Fluidigm BioMark microfluidic assay. In all nasal mucosa of non-infected animals, IFNβ mRNA was undetectable and comparisons for this gene were performed against levels of expression found in animals infected on 1 dpi. For IFNλ3 very low basal levels, at the limit of detection (Cq=25), were found in 2 out of 3 animals on 0 dpi. All other gene transcripts were detected at basal levels in control non-infected animals and were used as calibrator values (B in S3 Table).

At the level of MERS-CoV infected nasal epithelia (positively stained by IHC), most of the transcription variations occurred and peaked at 2 dpi. Genes coding for IFNβ (mean of 200 Fc) and IFNλ3 (mean of 350 Fc), and to a lesser extent IFNα (mean of 6 Fc) and IFNλ1 (mean of 11 Fc) were significantly upregulated. Relative expression of type I IFNs (α and β) and type III IFNs (λ1 and λ3) decreased progressively on 3 and 4 dpi (Fig 3A and B).

**Fig 3.**
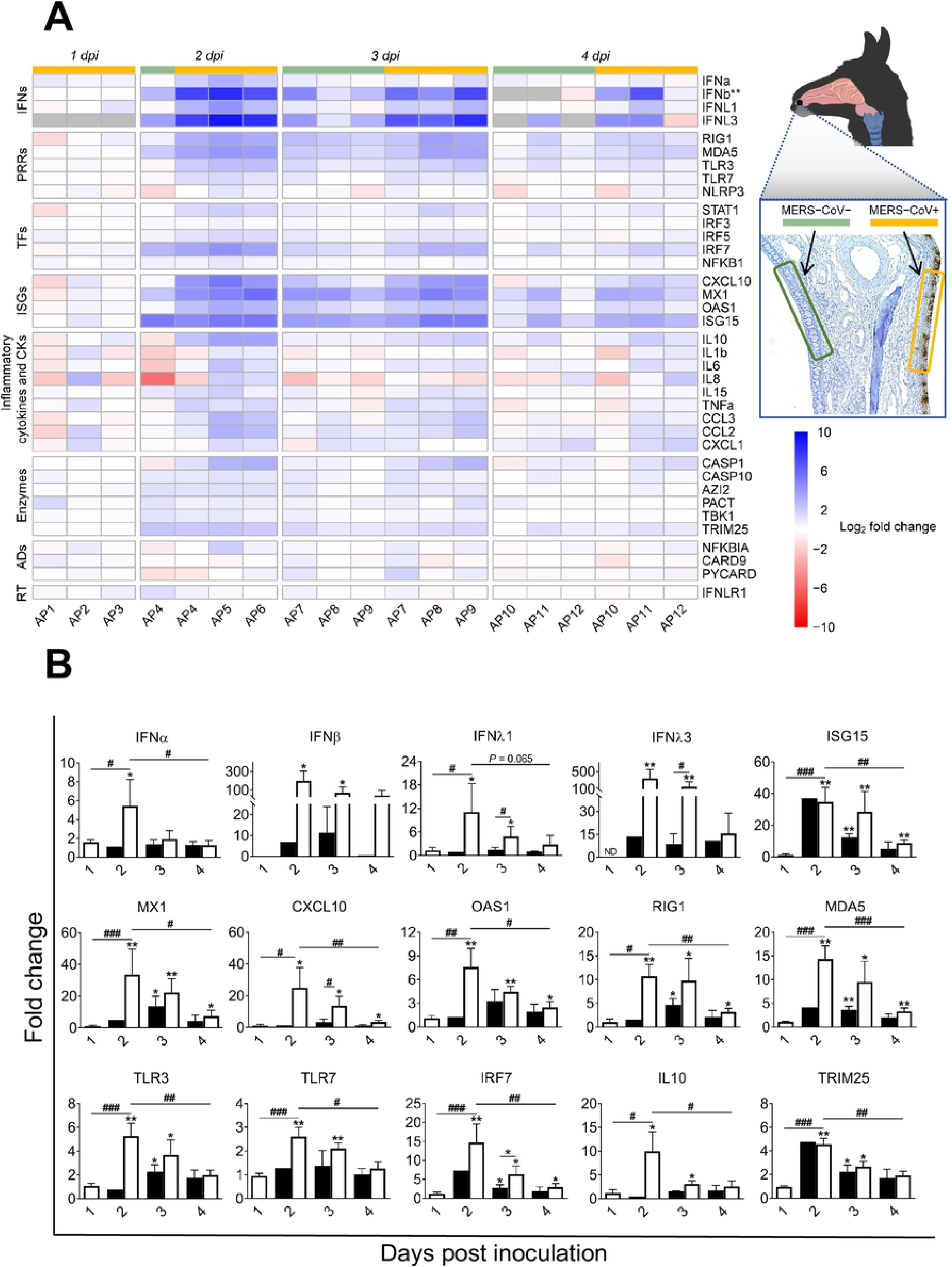
Kinetics of innate immune response genes induced at the nasal epithelia from MERS-CoV Qatar15/2015 infected alpacas. (A) The nasal epithelium of each alpaca (AP1 to AP15) was micro-dissected, as assessed by IHC (see the image of the nasal turbinate section in the upper right). Infected (orange)/non-infected (green) areas were isolated by LCM for RNA extraction and conversion to cDNA. The Fluidigm Biomark microfluidic assay was used to quantify transcripts of innate immune genes at different dpi. After normalization, Fc values between controls and infected animals were calculated. The resulting heatmap shows color variations corresponding to log2 Fc values; blue for increased and red for decreased gene expression values of infected animals compared to control animals, respectively. IFNβ was normalized with 1 dpi samples because it was not detected at 0 dpi. Grey rectangles indicate no detection of the corresponding gene. TFs, transcription factors; CKs, chemokines; ADs, adaptors; RT, receptor. MERS-CoV+ and MERS-CoV-, MERS-CoV positive and negative epithelium areas as assessed by IHC, respectively. (B) Average Fc of IFNs, ISGs, PRRs, IRF7, IL10 and TRIM25 genes in MERS-CoV+ nasal epithelia (white bars) and MERS-CoV-nasal epithelia (black bars) as assessed by IHC. Data are shown as means of ±SD. Statistical significance was determined by Student’s *t*-test. *\*P* < 0.05; *\*\*P* < 0.01; *\*\*\*P* < 0.001 when compared with the average values of non-infected alpacas (n = 3); *#P* < 0.05; *##P* < 0.01; *###P* < 0.001 when comparisons between groups are performed at different dpi.

The same patterns of gene transcription were observed for ISGs with antiviral activity (ISG15, MX1, CXCL10 and OAS1), IRF7, cytoplasmic viral RNA sensors (RIG1 and MDA5) and, although moderately upregulated, the endosome viral double and single stranded RNA sensors (TLR3 and TLR7 respectively) and the transcription factor STAT1. The E3 ubiquitin ligase TRIM25, an important enzyme that plays a key role in RIG1 ubiquitination, was upregulated on 2 dpi (mean of 4.5 Fc) and decreased afterward (Fig 3A and B, and C in S3 Table).

The pro-inflammatory antagonist IL10 was upregulated at 2 dpi (mean of 10 Fc) and returned to nearly steady state levels in the following days (Fig 3A and B). Only three genes involved in inflammation, TNFα, IL6 and IRF5 (mean of 2.5-3 Fc for each of them), were found slightly upregulated at 2 dpi, while the levels of expression of other critical pro-inflammatory factors, such as the cytokines IL8, IL1β, the adaptor PYCARD and the PRRs NLRP3, were marginally or not affected by the infection (Fig 3A, A in S3 Fig and C in S3 Table). Although not statistically significant, CASP1, an essential component of the inflammasome (as PYCARD and NLRP3), experienced moderate mRNA increases (2 to 8 Fc following the animals) at 2 and 3 dpi (A in S3 Fig). Finally, the chemo-attractant chemokines CCL2 and CCL3 were non-significantly upregulated (mean of 5 Fc) as shown in A in S3 Fig.

Nasal epithelium negative by IHC for MERS-CoV exhibited a moderate but nonsignificant increase for type I and III IFNs mRNA on 2, 3 and 4 dpi. However, ISG15, MX1, RIG1, MDA5, TLR3, IRF7, IL10 and TRIM25 genes were upregulated at 3 dpi but to a lower degree than in positive IHC epithelial areas (Fig 3A and B). Invariably, all the above-mentioned genes had a significant decreased expression from 2 dpi onwards. Transcription of most of the inflammatory cytokines was not affected or slightly downregulated upon infection (Fig 3A, A in S3 Fig and C in S3 Table).

As shown in S4 Fig, increased induction of type I IFNs in nasal epithelial micro-dissected samples was weakly correlated to increased viral MERS-CoV loads and transcription levels of IL10. In contrast, upregulation of type III IFNs had a stronger correlation with higher viral loads, as assessed by microfluidic PCR quantification of the viral M mRNA (r = 0.64, *P* = 0.004 for IFNλ1 and r = 0.64, *P* = 0.0163 for IFNλ3), the UpE gene (r = 0.65, *P* = 0.0033 for IFNλ1 and r = 0.64, *P* = 0.0163 for IFNλ3) and increased relative mRNA levels of IL10 (r = 0.9, *P* <0.0001 for IFNλ1 and r = 0.8, *P* = 0.0009 for IFNλ3). Our data indicate that antiviral mechanisms are set during the peak of infection in the nasal epithelia with a timely induction of type I and III IFNs, ISGs and IL10 without exacerbation of pro-inflammatory cytokines.

### ISGs and PRRs but not IFNs are upregulated in the nasal submucosa of infected alpacas: evidence for a potential IFN paracrine signaling

To further investigate the repercussion of MERS-CoV infection in the mucosa, innate immune responses occurring in the underlying submucosa were determined. Microdissected areas from the submucosa of control and infected animals (1 and 3 dpi) were analyzed for relative mRNA expression of innate immune genes. Except for IFNλ3 which was not detected in any inoculated or control animals, all other IFNs were expressed at basal levels in control non-infected alpacas. IFNβ was undetected at 1 dpi in two inoculated animals (AP2 and AP3) and below basal levels in alpaca AP1 while IFNα and IFNλ1 were marginally fluctuating. Most of the major transcriptional modifications occurred at 3 dpi. In submucosa underlying the most infected epithelia areas, IFNλ1 was expressed at basal levels while IFNα and IFNβ were not induced. On the contrary, ISGs, IRFs and PRRs, like ISG15 (mean of 77 Fc), OAS1 (mean of 6 Fc), MX1 (mean of 46 Fc), CXCL10 (mean of 14 Fc), RIG1 (mean of 12 Fc), MDA5 (mean of 16 Fc), IRF7 (mean of 9 Fc) and TLR7 (mean of 5 Fc) were highly to moderately upregulated. Expression of genes constituting the inflammasome was mostly not altered for NLRP3 or moderately increased for CASP1 (mean of 5 Fc) and PYCARD (mean of 3 Fc); but remarkably, IL1β was significantly downregulated (mean of 3 Fc; Fig 4A and B). Also, except for a non-significant increase of TNFα (mean of 5 Fc) and a slight upregulation of IRF5 (mean of 3 Fc; *P* < 0.05), no other pro-inflammatory factors were upregulated. Indeed, IL8 was downregulated (mean of −40 Fc). According to these findings, the antiinflammatory cytokine IL10 (mean of 3.5 Fc) was moderately upregulated (Fig 4A, B and D in S3 Table). Furthermore, the submucosa underlying epithelial cells with no MERS-CoV IHC labelling showed the same patterns of cytokine profiles as for that underlying heavily MERS-CoV infected mucosa but to a lesser degree of expression (Fig 4A and B). Overall, results obtained in the submucosa are indicative of a potential IFN paracrine signaling from the epithelium for the occurrence of ISGs induction in the absence of significant IFN mRNA synthesis. Moreover, despite mild focal infiltrations (Fig 2), genes involved in the inflammatory process were either suppressed, unaltered or mildly activated.

**Fig 4.**
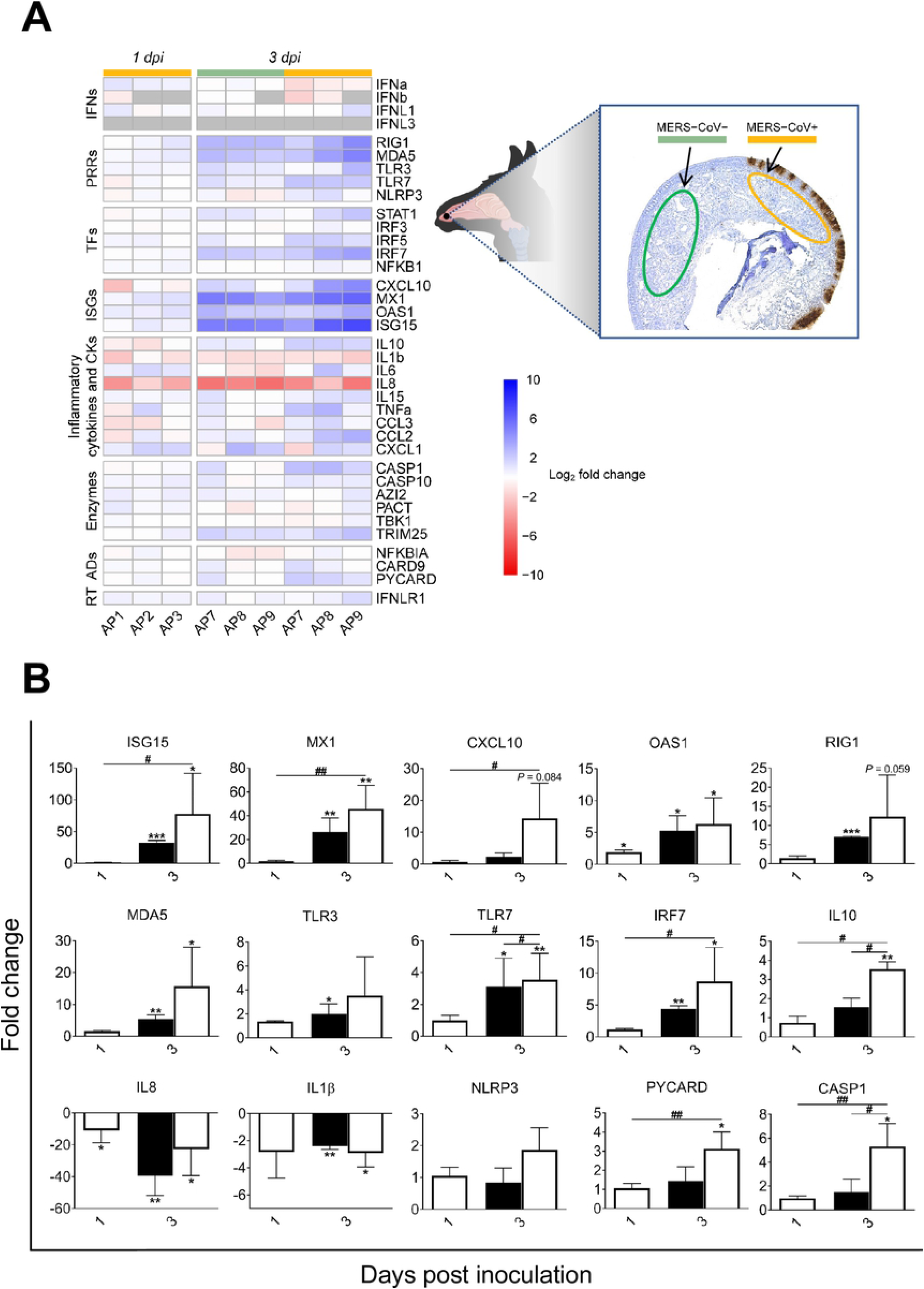
Kinetics of alpaca innate immune responses at the nasal submucosa in response to MERS-CoV Qatar15/2015. (A) The nasal submucosa of each alpaca (AP1, 2, 3, 7, 8, 9, 13, 14 and 15) was micro-dissected and areas underlying infected (orange)/non-infected (green) epithelium, as assessed by IHC (see the right panel), were selected and isolated for RNA extraction and conversion to cDNA. The Fluidigm Biomark microfluidic assay was used to quantify transcripts of innate immune genes at different dpi. After normalization, Fc values between controls and infected animals were calculated. The resulting heatmap shows color variations corresponding to log2 Fc values; blue for increased and red for decreased gene expression, respectively. The grey rectangles indicate no detection of the corresponding gene. TFs, transcription factors; CKs, chemokines; ADs, adaptors; RT, receptor. MERS-CoV+ and MERS-CoV-, MERS-CoV positive and negative epithelium areas as assessed by IHC, respectively. (B) Average Fc of ISGs, PRRs, IRF7, IL10, IL1β, IL8, NLRP3, CASP1and PYCARD genes in the submucosa underneath IHC MERS-CoV+ nasal epithelium (white bars) and IHC MERS-CoV-nasal epithelium (black bars). Data are shown as means of ±SD. Statistical significance was determined by Student’s *t*-test. \**P* < 0.05; \*\**P* < 0.01; \*\*\**P* < 0.001 when compared with the average values of non-infected alpacas (n = 3); *#P* < 0.05; *##P* < 0.01; ###*P* < 0.001 when comparisons between groups are performed at different dpi.

### ISGs are upregulated in the trachea of infected alpacas without increased endogenous IFNs mRNA

Innate immune response genes were further monitored along the URT. In trachea, IFNβ and IFNλ3 mRNA transcripts were not detected at any time post-inoculation including on 0 dpi. From 1 to 4 dpi, expression of IFNα and λ1 was slightly downregulated or unaltered, compared to basal levels found in non-infected controls (Fig 5A and E in S3 Table). Genes coding for ISGs (ISG15, MX1 and OAS1) and PRRs (RIG1, MDA5 and TLR3) showed a significant mRNA upregulation mainly at 2 and 3 dpi. When compared with animals necropsied at 1 dpi, IRF7 was significantly upregulated (mean of 9 Fc) at 2 dpi (Fig 5A and B). The expression of IL10 was not significantly altered (Fig 5A and B in S3 Fig). The induction of proinflammatory cytokine genes was inhibited or remained unchanged at all post-inoculation points (Fig 5A, B and B in S3 Fig), except for AP6. This animal unlike other alpacas sacrificed at 2 dpi experienced an upregulation of proinflammatory cytokines and the inflammasome in the trachea. However, IL6 and PYCARD mRNA levels were almost unaltered (Fig 5A, B in S3 Fig and E in S3 Table). Excluding OAS1, AP6 was the animal with the highest expression levels of ISGs, RNA sensors, transcriptional factors and chemokines as shown in Fig 5A, B, D, B in S3 Fig and E in S3 Table. In that respect, AP6 could be considered as an out layer since, unlike other animals, it presented nasal discharges with the highest number of infected cells in nose and trachea and the highest viral loads in trachea (S1 Table and A in S3 Table). Moreover, when compared to AP4 and AP5, which displayed nearly basal levels of CCL2 and CCL3 transcripts (Fig 5C, A in S5 Fig and E in S3 Table), the tracheal submucosa of AP6 was moderately infiltrated by macrophages and lymphocytes (Fig 5D). This also provides indirect proof that the chemokines CCL2 and CCL3 were produced at the protein level and exerted chemoattraction.

**Fig 5.**
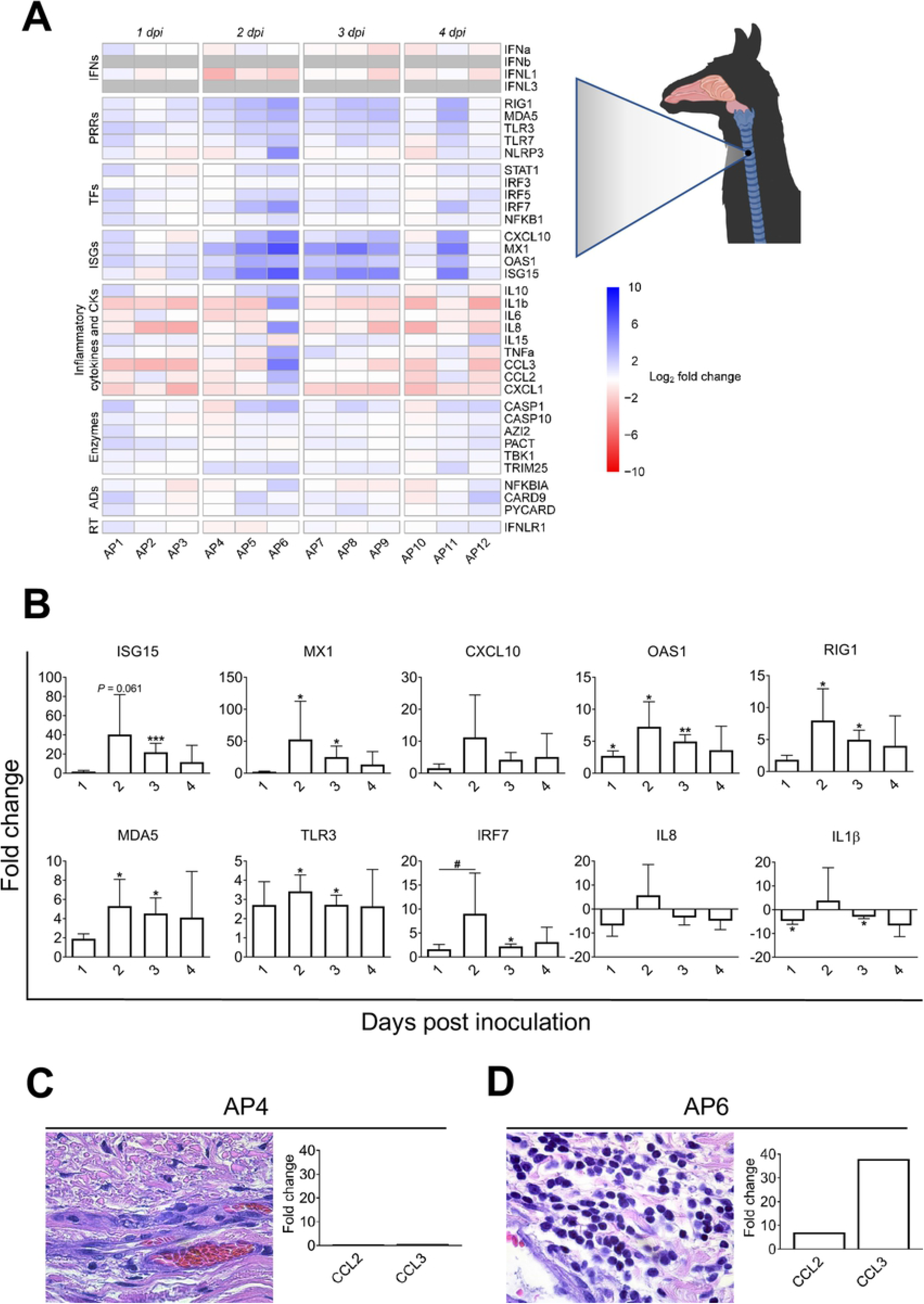
Kinetics of alpaca innate immune responses at the tracheal level in response to MERS-CoV Qatar15/2015. (A) Trachea samples were obtained by scraping MFPE sections from control (AP13-15) and infected alpacas (AP1-12), followed by RNA extraction and conversion to cDNA. A Fluidigm Biomark microfluidic assay was used to quantify transcripts of innate immune genes at different dpi. After normalization, Fc values between controls and infected animals were calculated. The resulting heatmap shows color variations corresponding to log2 Fc values; blue for increased and red for decreased gene expression, respectively. The grey rectangles indicate no detection of the corresponding gene. TFs, transcription factors; CKs, chemokines; ADs, adaptors; RT, receptor. (B) Average fold changes of ISGs, PRRs, IL8, IL1β and IRF7 genes in trachea. Data are shown as means of ±SD. Statistical significance was determined by Student’s *t-* test. \**P* < 0.05; \*\**P* < 0.01; \*\*\**P* < 0.001 (n = 3) when compared with non-infected alpacas (n = 3). (C) On 2 dpi, no inflammatory cells were observed in the tracheal submucosa of AP4 (see H/E stained MFPE tracheal section at the left). The graph at the right depicts Fc of mRNA transcripts for CCL2 and CCL3 in relation to control animals. (D) Infiltration of monocytes and lymphocytes in the tracheal submucosa of AP6 (see H/E stained MFPE tracheal section at the left) was associated with increased mRNA transcripts of CCL2 and CCL3 (see the bar graph at the right).

### Early induction of CCL2 and CCL3 correlates with infiltration of macrophages and lymphocyte like cells in the lungs of infected alpacas

Despite low infectivity of MERS-CoV in the LRT, infiltration of leukocytes was observed suggesting that cytokines and chemokines are at play. At the lung level, IFNα and IFNλ1 were expressed on 0 dpi in non-infected controls. In response to MERS-CoV, transcripts of IFNα and IFNλ1 fluctuated around basal levels (Fig 6A and F in S3 Table). IFNλ3 and IFNβ mRNAs were not detected in any of the animals including controls. Upregulation of ISGs (ISG15, MX1 and OAS1) and IRF7 started at 1 dpi, reached a peak at 2 dpi and weaned at 4 dpi. RIG1 and MDA5 were unevenly upregulated with a lower intensity than for other tissues (C in S3 Fig and F in S3 Table). Remarkably, NLRP3 was moderately but significantly upregulated at 1, 2 and 4 dpi concomitant with increased levels of TNFα and IL1β at 1 dpi (Fig 6A and B). However, transcription of CASP1 remained unaltered and the PYCARD gene was downregulated at 1, 2 and 4 dpi or unaltered at 3 dpi (C in S3 Fig and F in S3 Table). IL15, a natural killer cell activator cytokine, was significantly upregulated at 1 and 2 dpi while expression patterns of other proinflammatory cytokines varied between animals and days of infection (Fig 6A and B, C in S3 Fig and F in S3 Table).

**Fig 6.**
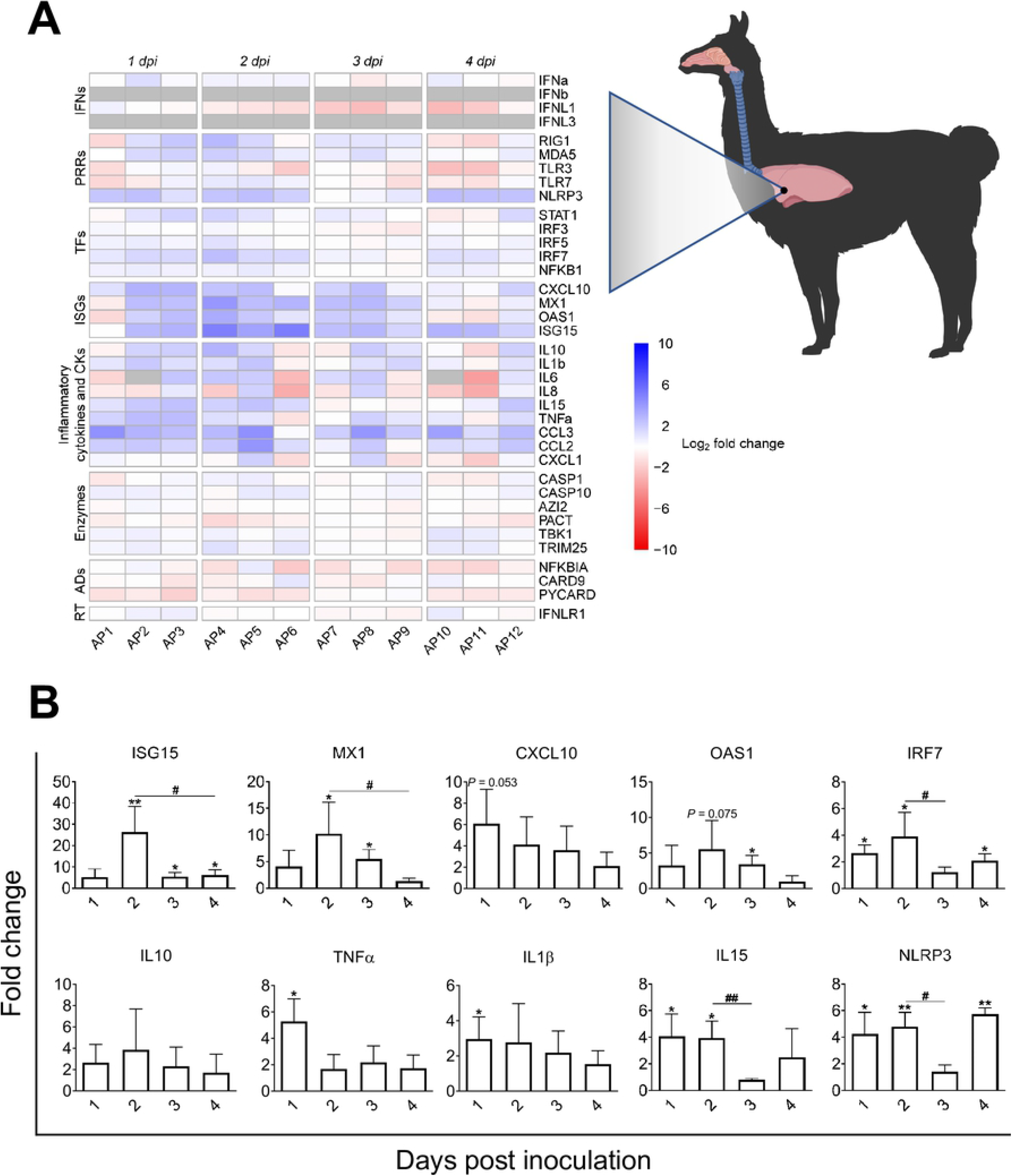
Kinetics of alpaca innate immune responses in the lungs following MERS-CoV Qatar15/2015 infection. (A) The lung sample of each infected alpaca (AP1 to AP15) was isolated for RNA extraction and conversion to cDNA. A Fluidigm Biomark microfluidic assay was used to quantify transcripts of innate immune genes at different dpi. After normalization, Fc values between controls and infected animals were calculated. The resulting heatmap shows color variations corresponding to log2 Fc values; blue for increased and red for decreased gene expression, respectively. The grey rectangles indicate no expression of the corresponding gene. TFs, transcription factors; CKs, chemokines; ADs, adaptors; RT, receptor. (B) Bar graphs representing average mRNA fold changes of ISGs (ISG15, MX1, CXCL10 and OAS1), IRF7, NLRP3, the inflammatory cytokines IL1β, TNFα, IL10 and IL15 genes in lung. Data are shown as means of ±SD. Statistical significance was determined by Student’s *t*-test. \**P* < 0.05; \*\**P* < 0.01; \*\*\**P* < 0.001 (n = 3) compared with non-infected alpacas (n = 3); #*P* < 0.05; ##*P* < 0.01; ###*P* < 0.001 (n = 3) when comparisons between groups are performed at different dpi.

In addition, CCL2 and CCL3 were moderately to highly upregulated at most of the time points (Fig 7A, C in S3 Fig and B in S5 Fig). All animals showed very mild infiltration of leukocytes in the alveoli in response to MERS-CoV (Fig 7B and C in S5 Fig). Remarkably, AP6 showed accumulation of leukocytes without a notable induction of CCL2 and CCL3 mRNAs. Moreover, this animal was the highest producer of these chemoattractant chemokines in trachea. Nonetheless, the number of infiltrating leukocytes correlated with the induction of CCL2 (r = 0.62, *P* = 0.0236) and CCL3 (r = 0.59, *P* = 0.0323) as shown in D and E in S5 Fig. When considering AP6 as an out layer, correlations became stronger for both CCL2 (r = 0.72, *P* = 0.0077) and CCL3 (r = 0.87, *P* = 0.0002) as shown in Fig 7A. Overall, ISGs were induced in the absence of IFN gene expression suggesting an endocrine effect of IFNs produced in the nasal epithelia. In addition, despite transient accumulation of leukocytes correlating with induction of CCL2 and CCL3 in the lung, exacerbation of inflammatory responses did not occur.

**Fig 7.**
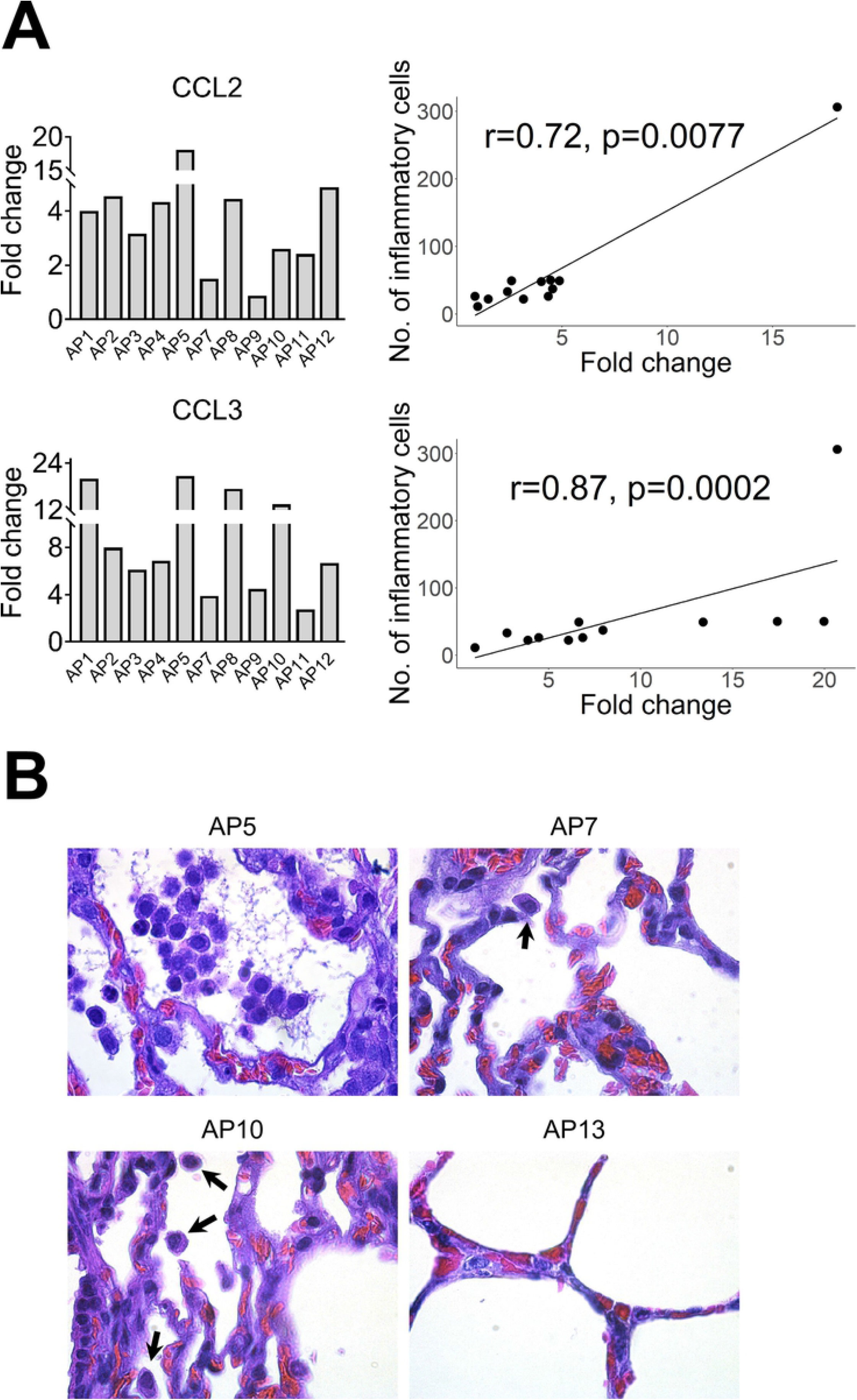
Infiltration of leukocytes in alveoli are correlated with upregulation of CCL2 and CCL3 mRNA. (A) Relative expression of CCL2 and CCL3 mRNA (Bar graphs at the left) was performed with a microfluidic PCR assay in the lung of alpacas (except AP6, see text). Leukocytes were counted in 3 microscopic fields (400X) per lung section in all animals (except AP6), including non-infected controls. Relative expression levels of CCL2 and CCL3 were plotted against the number of inflammatory cells (dot plot at the right). Correlation coefficients were established using the Spearman’s correlation test (right panel). (B) The number of leukocytes in alveoli was the highest in AP5, lower in AP10 (arrows), occasional in AP7 (arrow) and hardly detectable in AP13. Original magnification: ×1000 for all samples.

### Genes of the NF-κ B pathway and IRF5 are transcriptionally unaltered in respiratory tissues during MERS-CoV infection

Transcription of the NF-κB p50 subunit (NFKB1) and some important regulators of the NF-κB pathway were checked along URTs and LRTs. These regulators included two enzymes, AZI2 and TBK1, downstream of the RLRs and TLR signaling pathways, essential for the phosphorylation of NF-κB, the NF-κB inhibitor NFKBIA and an adaptor CARD9 that mediates signals from C-type lectins to regulate the NF-κB pathway (S6 Fig). All these genes were expressed at basal levels in the nasal mucosa, submucosa, trachea and lung of control non-infected alpacas. NFKB1, AZI2 and TBK1 were only marginally induced or remained unaltered in the nasal epithelia; however, NFKBIA and CARD9 were expressed at basal levels or even slightly downregulated in all the tissues. All the above mentioned genes were fluctuating around basal transcriptional levels across alpaca respiratory tracts (Fig 3A-6A and C-F in S3 Table) indicating that the NF-κB mediated signaling pathway was not exacerbated or downregulated in response to MERS-CoV. Moreover, IRF5, an inducer of TNFα and IL6 as well as an important factor for the polarization of M1 macrophages, was only slightly but significantly upregulated in the nasal epithelium (2 dpi) and submucosa (3 dpi). In all other tissues and, in particular lung, the majority of animals did not reach the threshold of 2 Fc for IRF5 (C in S3 Fig and F in S3 Table).

## Discussion

The present study exposes an integrated investigation at the respiratory tract level performed *in vivo* in a natural reservoir/intermediate host upon MERS-CoV infection. As camelids are the primary zoonotic reservoirs for human infection, it is essential to gain insight into the potential mechanisms of asymptomatic manifestation since they can unravel targets for prophylactic treatments in susceptible hosts. In the present study, all experimentally inoculated alpacas were successfully infected, as genomic viral UpE, subgenomic M mRNA and the N protein could be detected in most of the tissues examined by IHC, RT-qPCR or micro fluidic PCR assays from the URT and LRT at 1 to 4 dpi. This proved that the MERS-CoV Qatar15/2015 strain had spread and in some instance replicated throughout the whole respiratory tracts, although mainly at the URT where the virus receptor DPP4 (dipeptidyl peptidase-4) is most abundant [34]. Acknowledging that respiratory mucosal epithelial cells are the main primary target for MERS-CoV and the paucity of data on the effect *in vivo* of the virus to the mucosal barrier on humans and other laboratory species (mainly human DPP4 transgenic or transduced mice and non-human primates) a special attention was given here to monitor innate immune responses occurring during the first 4 days of infection in the nasal epithelium of alpacas and the possible repercussions of these responses in the respiratory tract.

In mucosa, pathogens are sensed by PRRs leading to the production of type I and III IFNs via IRFs. In turn, IFNs can activate through the JAK-STAT pathway, in an autocrine, paracrine or endocrine manner, transcription of various antiviral and regulatory ISGs including also PRRs and IRFs. Fine tuning of these responses is paramount in determining the outcome of viral infectious diseases [31]. In MERS-CoV infected alpacas, during the first 24h, apart from IFNβ which started to be induced, type I and III IFNs were not altered in the nasal mucosa when compared to control animals. This short period seems to be crucial for the virus to establish a productive infection in alpacas for at least 5 to 6 days [5] by delaying host protective mechanisms such as activation of PRR pathways [17,30,35]. Nonetheless, and only in the nasal epithelium, transcription of all type I and III IFNs peaked simultaneously at 2 dpi and weaned progressively. This wave of IFNs was concomitant with the induction of ISGs, PRRs and IRF7 not only at the nasal epithelial barrier and its underlying submucosa but also in trachea and lungs where IFN mRNAs were never detected or induced. Therefore, the effect of the IFN response at the nasal mucosa may extend to distant respiratory tissues in a paracrine or endocrine manner. In that respect, in mice, IFNλ produced in the URT prevents dissemination of influenza virus to the lung [36].

The use of IHC and microfluidic PCR assays combined with LCM permitted to compare, in the same and different animals, areas of the nasal epithelium with high and low viral loads. Heavily infected mucosal areas were responsible for the highest induction of type III IFNs. Although markedly induced, type I IFNs did not significantly correlated with tissue viral loads. Consistent with this IFN production, the number of MERS-CoV infected cells detected by IHC was maximal on 2 dpi and decreased over time. However, all infected alpacas shed infectious viral particles in the nasal cavity from 1 dpi onwards according to previous studies [5,6]. This might be due to a high accumulation of virus in the nasal cavity which may take a few days to clear. Remarkably, results obtained herein contrast with those found in human epithelial cells and respiratory explants since the induction of type I and III IFNs was limited or inhibited in response to MERS-CoV [19,20,23]. The dampened IFN priming in humans is likely due to the role of IFN antagonism by MERS-CoV accessory proteins [20,24,29,37], through which virus replication is facilitated. It is important to note that the MERS-CoV strain (Qatar15/2015) used herein is from human origin and, unlike some dromedary isolated African strains, it does not display any deletions in the above-mentioned non-structural proteins [22]. Therefore, alpacas are able to overcome IFN inhibitory mechanisms in a relative short time with the induction of antiviral or IFN positive regulatory ISGs. Nonetheless, the prominent manifestation of MERS in humans occurs in lungs with a massive inflammation provoked by infiltration of macrophages and lymphocytes [38]. To the contrary, and although AP6 experienced discrete nasal discharges and displayed large numbers of infected nasal epithelial cells, alpacas only exhibited transient mild and focal infiltration of lymphocytes, macrophages and neutrophils in response to MERS-CoV. Such infiltrations were mainly observed on 2 dpi and gradually resolved afterwards. These findings suggested a rapid fine tuning of specific mechanisms to control inflammation mediated essentially by the mucosal barrier, leading to an efficient innate immune response. Indeed, type III IFNs are known to induce milder inflammatory responses than IFNß [31] and IL10 is an anti-inflammatory cytokine [39]. Furthermore, recent studies in mice indicate intricate mechanisms between IFNλ and IL10 via dendritic cells (DC) conditioning protective mucosal immunity against influenza virus [40]. In this view, it is noteworthy to mention that plasmacytoid DC are seemingly the only cell-type that produce *in vitro* type I and III IFNs after MERS-CoV infection in humans [41]. By contrast, *in vivo,* pDCs which predominantly localize in the submucosa are not the main IFN producers [42,43]. Due to the lack of specific cell markers for DC in alpacas, detection of these cells in tissues and their contribution to innate immunity could not be performed. Nonetheless, a strong positive correlation between IFNλ1,3 and IL10 mRNA levels was observed in the nasal epithelium of alpacas and this could reflect a physiological cross-regulation between these structurally related cytokines upon MERS-CoV infection. By contrast, the weak correlation between IFNβ and IL10 mRNA induction suggests that only type III IFNs are directly or indirectly mediating antiinflammatory action. Furthermore, IL10 was significantly up regulated, although at moderate levels, in the nasal submucosa underlying heavily infected epithelial cells. Moreover, IRF5 known to be a pro-inflammatory transcription factor and a key element in the M1 polarization of macrophages [44,45] was only slightly induced in the nasal mucosa and submucosa on 2 and 3 dpi. However, no IRF5 mRNA increased in the lung despite infiltration of mainly macrophages and lymphocytes as early as 1 dpi. Accordingly, CCL2, CCL3, CXCL10 and TNFα were induced in the lung in the absence of type I IFNs, as found for MERS-CoV infected human monocyte-derived macrophages [16]. With the observation that other pro-inflammatory cytokines are either weakly or not induced, it is suspected that inflammatory M1 macrophages are not abundant in the lung of infected alpacas. Also, IL15, an activator and attractor of NK cells and macrophages [46], was only induced in lung as soon as 1 dpi, suggesting a role for clearance of the virus by these cells. Strikingly, NLRP3 mRNA was more abundant in the lung (with the exception of AP6 in trachea), maybe due to the presence of infiltrating macrophages where NLRP3 is mostly produced [47,48]. However, CASP1 and PYCARD, the two other components of inflammasome, were not induced. This scenario, combined with a discrete upregulation of IL1β and IL6 in some animals, will not probably result in a cytokine storm in the lung. Accordingly, in other respiratory tissues, the major proinflammatory cytokines were occasionally mildly induced, suggesting also a functional but attenuated NF-κB signaling [49]. Moreover, during infection, genes involved in the regulation of the NF-κB cascade (AZI2, TBK1 and NFKBIA) and NFKB1were transcriptionally unaltered in all the respiratory tract of alpacas preventing acute inflammation in response to MERS-CoV.

In a mice model of MERS-CoV [35], IFNs production and MERS-CoV replication peaked simultaneously in lungs at 2 dpi resulting in a sublethal infection. While early induction or treatment with IFNß were crucial for virus clearance and induction of protective immunity, late administration of IFNs led in lungs to enhanced infiltrations of monocytes, macrophages, neutrophils, expression of proinflammatory cytokines and increased mortality. Also, indirect proof was given that IFNs synthesis was under sensing of TLR7 but not MAVS in airway epithelial cells. However, this study did not report the effect of MERS-CoV infection in the URT [35]. Benefits of early type I IFN treatment were also shown for non-human primates [50] but IFN therapy on humans has failed so far certainly because it was applied essentially on critically ill patients [51]. Thus, some differences are noticed between a MERS-CoV sublethal infection in mice and that occurring in a natural host. Here, we highlighted the essential role of the nasal mucosa as a main producer of IFNs upon MERS-CoV infection driving in combination with IL10 a mild inflammatory response along the respiratory tract. The observation in alpacas of a dimmed NLRP3 inflammasome is consistent with that found in bat primary immune cells infected in vitro with MERS-CoV [32], suggesting a primary role in the control of inflammation in reservoir/intermediary hosts despite high viral loads. Further, this study provides mechanistic insights on how innate immunity overcomes a MERS-CoV infection. A proposed model of cytokine and signaling pathways interactions resuming innate immune responses upon MERS-CoV infection in the respiratory tract of alpacas is depicted in Fig 8. Inherent to any study performed with large outbred animals under BSL3 containment, the present study suffers some limitations. First, the number of animals used was kept to minimal to avoid prolonged exposure of personnel during sampling and necropsies hindering some statistical significances with variations between animals. Second, the LCM and microfluidic PCR allowed determination of gene expression at the tissue but not the single cell level. This combined with the lack of antibodies defining cell types in alpaca and working in IHC impeded further characterization of DCs and leukocytes in tissues.

**Fig 8.**
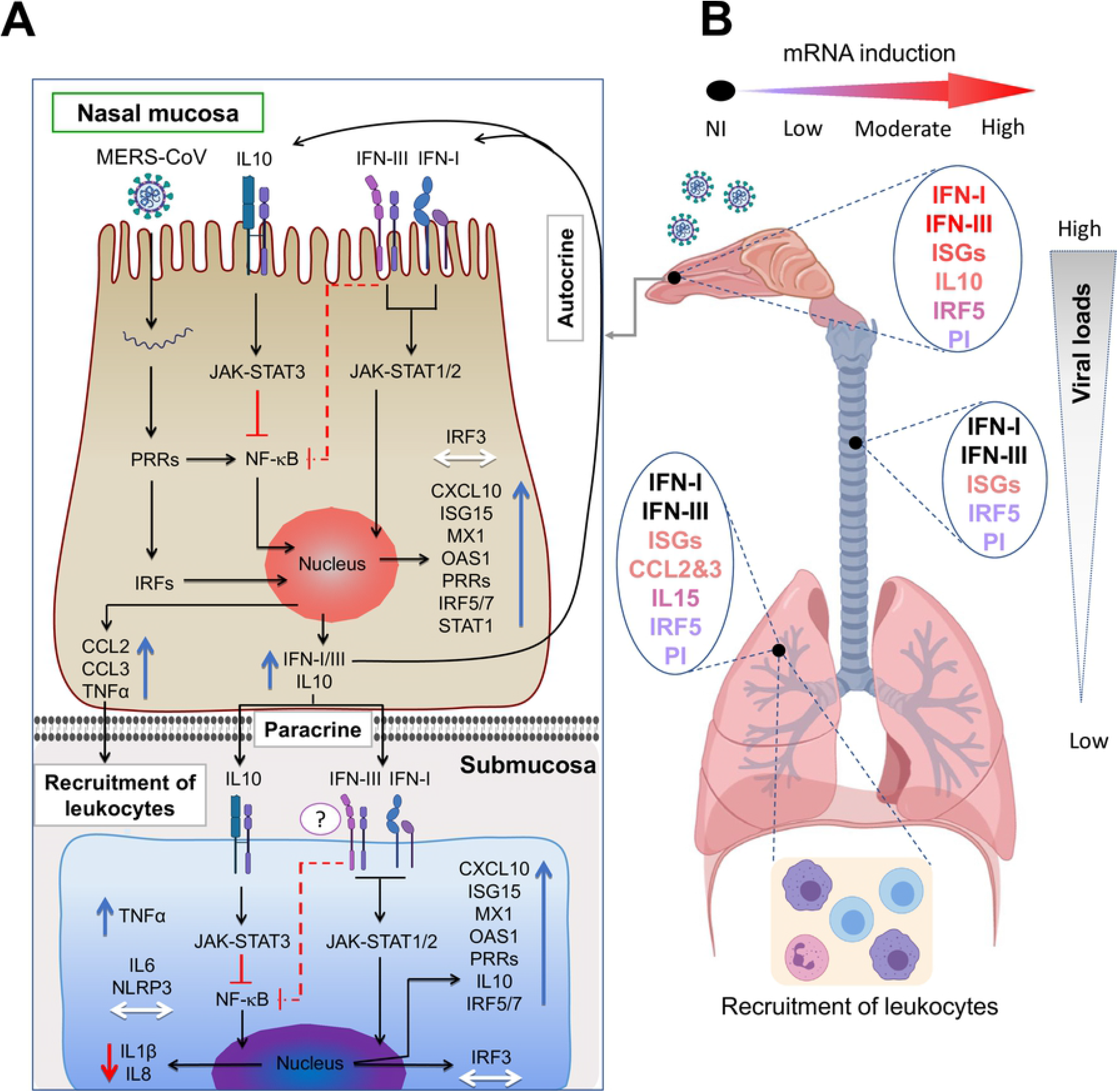
Proposed mechanistic model for MERS-CoV-induced protective innate immune responses in alpaca respiratory tracts. (A) Upon MERS-CoV infection of nasal epithelial cells, PRRs and IRFs are engaged to induce IFNs and ISGs exerting their effects in an autocrine or paracrine manner. Upregulation of IL10 combined with the action of type III IFNs will result in a dimmed synthesis of proinflammatory cytokines under NF-κB control. Induced chemokines lead to focal, mild infiltration of leukocytes. The paracrine effect of IFNs is evidenced in the nasal submucosa where ISGs are upregulated without endogenous induction of IFNs. IL10 and type III IFNs will act on the NF-κB pathway preventing production of IL8 and IL1β mRNAs depriving the NLRP3 inflammasome of its substrate. Slight increased levels of IRF5 mRNAs will indicate the presence of few M1 macrophages in the submucosa. Blue, red and white arrows indicate upregulation, downregulation and unaltered gene transcription respectively. (B) Concomitant to decreased viral loads, transcription of ISGs is lowered in trachea and lungs where IFNs are not induced. Infiltration of mainly mononuclear leukocytes occurs in the lungs as a result of chemokine synthesis in the absence of IRF5 induction but upregulation of IL15. Thus, reduced number of M1 macrophages and activation of NK cells will contribute to a controlled inflammation and clearance of the virus. PI, Pro-inflammatory cytokines; NI, not induced.

## Material and methods

### Ethics statement

All animal experiments were approved by the Ethical and Animal Welfare Committee of IRTA (CEEA-IRTA) and by the Ethical Commission of Animal Experimentation of the Autonomous Government of Catalonia (file N° FUE-2018-00884575 – Project N°10370). The work with the infectious MERS-CoV Qatar15/2015 strain was performed under biosafety level-3 (BSL-3) facilities of the Biocontainment Unit of IRTA-CReSA in Barcelona, Spain.

### Cell culture and virus

A passage 2 of MERS-CoV Qatar15/2015 isolate (GenBank Accession MK280984) was propagated in Vero E6 cells and its infectious titer was calculated by determining the dilution that caused cytopathic effect (CPE) in 50% of the inoculated Vero E6 cultures (50% tissue culture infectious dose endpoint [TCID50]), as previously described [9].

### Laser capture microdissection (LCM)

For each animal, four consecutive sections from the same MFPE-block containing nasal specimens were cut and processed as described in the supplementary material and method section. One of the sections was subjected to IHC to localize infected/non infected cells in the tissues and served as a reference (template) for the three subsequent sections which were subjected to LCM. Infected and non-infected nasal epithelium (mucosa) areas, as assessed by IHC in the template section, as well as their respective underlying submucosa areas were delineated and micro-dissected using the Leica LMD6500 (Leica AS LMD; Wetzlar, Germany) system (6.3× magnification, Laser Microdissection 6500 software version 6.7.0.3754). More detailed information is provided in S1 Fig for visualization of the microdissection process.

Dissected specific areas from each three MFPE sections were measured and introduced into RNase-free 0.5 ml Eppendorf tubes with buffer PKD from the miRNeasy FFPE Kit (Qiagen, Valencia, CA, USA). Direct microdissection of IHC stained MFPE sections were attempted but failed to provide RNA with enough quality and yields for further microfluidic quantitative PCR assays.

### Total RNA isolation and cDNA synthesis

Total RNA extraction from micro-dissected or scraped tissues was performed using the miRNeasy FFPE Kit, following the manufacturer’s instruction. Isolated RNA was concentrated by precipitation in ethanol, purified using RNeasy MinElute spin columns (Qiagen, Hilden, Germany) and treated for 10 min with DNase I (ArcticZymes, Norway). cDNA was generated from 110 ng of total RNA using the PrimeScript™ RT reagent Kit (Takara, Japan) with a combination of both oligo-dT and random hexamers following manufacturer’s instructions. Additionally, total RNA extracts from MERS-CoV infected nasal epithelia of AP5, 6, 7, 8, 9 and 11 were pooled at the same proportion per animal and used to generate cDNA controls for validation of gene expression assays.

### Selection of innate immune genes, primers design and microfluidic quantitative PCR assay

Thirty-seven innate immune genes were selected to study the gene expression and transcriptional regulation of the main known canonical signaling pathways acting on antiviral innate immunity and inflammation (S6 Fig). Primer design and validation for an efficient cDNA amplification (S2 Table) is described in detail in the S1 Text section (S1 Text). cDNA from micro-dissected or scraped tissue samples were used to quantify camelid gene expression levels in duplicates by a microfluidic qPCR with the 96.96 Dynamic Array integrated fluidic circuit of the Biomark HD system (Fluidigm Corporation, South San Francisco, USA), according to the manufacturer’s instructions (see S1 Text).

### Relative quantification of innate immune response genes and statistical analyses

Data analyses and normalization of microfluidic qPCR assays is described in the S1 Text section. Briefly, the relative standard curve method (see Applied Biosystems User Bulletin #2 https://www.gu.se/digitalAssets/1125/1125331_ABI_Guide_Relative_Quantification_using_realtime_PCR.pdf) was applied to extrapolate the quantity values of targets and normalizer genes of the studied samples. Then, the target amount is normalized by using the combination of three reference control genes. The normalized quantity (NQ) values of individual samples from infected animals at different dpi were used for comparison against the NQ mean of three non-infected control animals (calibrator) per assay. Thus, the up- or down-regulated expression of each gene was expressed in fold changes (Fc) after dividing each individual NQ value with the calibrator. Unlike all other tested genes, IFNβ mRNA was undetectable in the nasal epithelia of non-infected control animals, therefore, comparisons for this gene in the nasal epithelia were performed in relation to the mean of IFNβ levels of three infected alpacas on 1 dpi. C-F in S3 Table recompiles all the results expressed in Fc relative to the control alpacas obtained with the Fluidigm microfluidic assay for all 37 genes tested. A in S3 Table shows the quantification, expressed in Cq values, of MERS-CoV genomic (UpE) and subgenomic RNA (M mRNA), obtained with the microfluidic qPCR assay. B in S3 Table indicates the expression levels (in Cq values) of all genes from non-infected alpacas euthanized at 0 dpi. The limit of detection for expressed genes was Cq=25. A logarithmic 10 transformations were applied on Fc values to approach a log normal distribution. Thus, the Student’s Ltest could be used to compare the means of the logarithmic Fc obtained at different dpi for each group of animals. Significant upregulation or downregulation of genes was considered if they met the criteria of a relative Fc of ?2-fold or ≤2-fold respectively with *P* < 0.05. Having determined that RT-qPCR data (Cq values) are normally distributed according to the Shapiro-Wilk normality test, the Tukey’s multiple comparisons test was applied to compare Cq values of tissue samples at different dpi. Differences were considered significant at *P* < 0.05. Correlation coefficients were determined using the Spearman’s correlation test.

## Acknowledgments

We thank Bart L. Haagmans from Erasmus Medical Center for providing the MERS-CoV Qatar15/2015 strain; Montse Amenós and Joana Ribes from the Centre for research in agricultural Genomics for guidance on the use of LCM and Fluidigm BioMark microfluidic assay. We thank Isabelle Schwartz from INRAe for critically reviewing the manuscript. We are particularly indebted with staff of the BSL3 bio containment animal facility at CReSA.

## Author Contributions

**Conceptualization:** Nigeer Te, Jordi Rodon, Maria Ballester, Mónica Pérez, Lola Pailler-García, Joaquim Segalés, Júlia Vergara-Alert, Albert Bensaid.

**Funding acquisition:** Joaquim Segalés, Júlia Vergara-Alert, Albert Bensaid.

**Investigation:** Nigeer Te, Jordi Rodon, Joaquim Segalés, Júlia Vergara-Alert, Albert Bensaid.

**Supervision:** Joaquim Segalés, Júlia Vergara-Alert, Albert Bensaid.

**Writing – original draft:** Nigeer Te, Jordi Rodon, Joaquim Segalés, Júlia Vergara-Alert, Albert Bensaid.

**Writing – review & editing:** Nigeer Te, Jordi Rodon, Maria Ballester, Joaquim Segalés, Júlia Vergara-Alert, Albert Bensaid.

## Supporting information

**S1 Text. Material and methods**

**S1 Fig. Laser capture microdissection (LCM) of the nasal turbinate mucosa and underlying submucosa of MERS-CoV infected alpacas.** For each animal, four consecutive 6-7 μm sections from the same methacarn-fixed paraffin-embedded (MFPE) block were performed and mounted onto Leica RNase-free PEN slides. One of the sections was stained by IHC to detect the presence of the viral N protein and visualize heavily infected mucosa areas (A) from non-stained areas apparently devoided of virus (E). This IHC stained section served as a template to localize by overlapping on the other contiguous MFPE sections, stained only with cresyl violet, infected (B) and “non-infected” (F) areas prior LCM. Then LCM was applied to collect the respective selected areas in the mucosa (C and G) and the underlying submucosa (D and H).

**S2 Fig. MERS-CoV UpE gene and M mRNA loads in MFPE samples.** Micro-dissected (Nasal epithelia and underlying submucosa) and scrapped (Trachea and lung) MFPE tissue sections were prepared on the basis of an overlapping template section stained by IHC to localize the MERS-CoV N protein as described in S1 Fig. RNA extracted from these samples collected in alpacas prior (0 dpi, n=3) MERS-CoV inoculation and after during 4 consecutive days (1 to 4 dpi, n=3 per day) were converted into cDNA and (A) the MERS-CoV UpE gene and (B) the M mRNA amplified with a PCR microfluidic assay (Fluidigm Biomark). Error bars indicate SDs when results were positive in more than one animal. At 1 dpi, only Epi+ were sampled by LCM, the majority of cells were IHC negative. At 2 dpi, only one animal displayed distinct Epi-areas within the nasal turbinates which could be micro-dissected. All other animals at 2 dpi had a massive infection of nasal turbinates with few IHC negative cells. A in S3 Table provides detailed information on Cq values obtained for each animal and indicates also the samples with no detectable viral RNA in the nasal submucosa trachea and lung. Abbreviations: Epi-, non-infected nasal epithelia, as assessed by IHC; Epi+, MERS-CoV infected nasal epithelia, as assessed by IHC; Sub, submucosa underlying non-infected nasal epithelia, as assessed by IHC; Sub+, submucosa underlying MERS-CoV infected nasal epithelia, as assessed by IHC; MFPE, methacarn-fixed embedded-tissues; LCM, laser capture microdissection.

**S3 Fig. Innate immune response genes induced at the nasal epithelia, trachea and lung from MERS-CoV Qatar15-2015 infected alpacas.** Micro-dissected nasal epithelia and scrapped (Trachea and lung) MFPE tissue sections were prepared on the basis of an overlapping template section stained by IHC to localize the MERS-CoV N protein as described in S1 Fig. RNA extracted from these samples collected in alpacas prior (0 dpi, n=3) MERS-CoV inoculation and after during 4 consecutive days (1 to 4 dpi, n=3 per day) were converted into cDNA. The Fluidigm Biomark microfluidic assay was used to amplify and quantify gene transcripts in (A) MERS-CoV positive nasal epithelia (white bars) and MERS-CoV negative nasal epithelia (black bars) as assessed by IHC (B), trachea and (C) lung at different dpi (1 to 4 dpi). At 1 dpi, only MERS-CoV positive nasal epithelia was sampled by LCM, the majority of cells were IHC negative. At 2 dpi, only one animal displayed distinct MERS-CoV negative areas within the nasal turbinates which could be micro-dissected. All other animals at 2 dpi had a massive infection of nasal turbinates with few IHC negative cells. Data are shown as means of Fc±SD. Statistical significance was determined by Student’s t-test. \**P* < 0.05; \*\**P* < 0.01; \*\*\**P* < 0.001 (n = 3) when compared with the average values of non-infected alpacas (n = 3); #*P* < 0.05; ##*P* < 0.01; ###*P* < 0.001 (n = 3) when comparisons between groups are performed at different dpi. Significant upregulation of genes was considered if they met the criteria of a relative fold change of ≥ 2-fold with *P* < 0.05; similarly, significant downregulation of genes was considered if they met the criteria of a relative fold change of ≤2-fold with *P* < 0.05.

**S4 Fig. MERS-CoV loads and IL10 upregulation correlate with induction of type III IFNs but not type I IFNs in microdissected nasal epithelia.** Portions of the nasal epithelia of each alpaca (AP1 to AP15) were micro-dissected from MFPE tissue sections on the basis of a section template stained by IHC allowing selection of areas positive or negative for the MERS-CoV N protein (S1 Fig). After RNA extraction and conversion to cDNA the Fluidigm Biomark microfluidic assay was used to amplify, among others, transcripts IFNα, IFNβ, IFNλ1, IFNλ3 and IL10 and the viral UpE gene and M mRNA from all nasal epithelia (negative and positive by IHC) collected at different dpi (0 to 4 dpi). Values are expressed, after normalization with the HPRT1, GAPDH and UbC gene transcripts, as fold changes of the expression of cytokines in infected animals versus control animals (AP13-15 sacrificed at 0 dpi) and inverted Cq for the viral UpE gene and M mRNA (A in S3 Table and S2 Fig). IFNβ (IFNb) was normalized with 1 dpi samples because it was not expressed at 0 dpi. Inverted Cq values of MERS-CoV M mRNA and UpE, and relative expression levels of IL10 were plotted against relative expression levels of (A) IFNα, (B) IFNβ, (C) IFNλ1 and (D) IFNλ3 in micro-dissected nasal epithelia. Correlation coefficients were established using the Spearman’s correlation test. dpi, days post inoculation.

**S5 Fig. Correlation between relative expression of chemokines and the number of inflammatory cells in alpaca trachea and lung.** (A) No inflammatory cells nor CCL2 and CCL3 mRNA induction was seen in the MFPE tracheal submucosa of AP13 (0 dpi) and AP5 (2 dpi). (B) Relative expression of CCL2 and CCL3 mRNA was performed with a Fluidigm Biomark microfluidic assay in the lung of alpacas (including AP6). Inflammatory cells were counted in 3 microscopic fields (400X) per MFPE lung section in all animals, including non-infected controls. (C) The number of inflammatory cells in alveoli was high in AP6, lower in AP4, 8 and 12 (arrows), occasional in AP2 and 3 (arrow). Original magnification: ×1000 for all samples. Relative expression levels of (D) CCL2 and (E) CCL3 were plotted against the number of inflammatory cells in lungs. Correlation coefficients were established using the Spearman’s correlation. MFPE: methacarn-fixed paraffin-embedded.

**S6 Fig. Pattern recognition receptors (PRRs), IFN signaling and pro-inflammatory cytokine pathway involving antiviral innate immunity induced by RNA viruses.** Upon sensing of RNA viruses, PRRs interact with their associated adapter proteins, thereby transmitting downstream signals to nuclear factor (NF-κB) and interferon regulatory factors (IRF3, IRF5 and IRF7) which further stimulate the production of pro-inflammatory cytokines (IL6, IL8, IL15, TNFα, CCL3, CCL2 CXCL1, NLRP3 and pro-IL1β) and IFNs (predominantly type I and III IFNs), respectively. Importantly, IRF5, is also a potent inductor of pro-inflammatory cytokines. The production of IFNs enhances the release of interferon stimulated genes (ISGs) which exert both, antiviral and host gene regulatory activities. IFNs can act in an autocrine and paracrine manner to induce the expression of ISGs via the JAK-STAT signaling pathway. While type I and III IFNs induce a similar set of ISGs (RIG1, MDA5, STAT1, IRF3, IRF5, IRF7, CXCL10, MX1, OAS1, ISG15, TRIM25), type I IFN signaling also activates pro-inflammatory cytokines and chemokines, which in turn recruit leukocytes to the site of infection. IL10 functions as an anti-inflammatory cytokine that activates STAT3 which is involved in the negative regulation of NF-κB. AZI2 and TBK1 (which also phosphorylate IRFs) participate positively in the NF-κB signaling pathway while IκBα inhibits translocation of NF-κB in the nucleus until degradation. Activation of the inflammasome, composed of NLRP3, PYCARD (ACS) and pro-CASP1 leads to the auto-cleavage of pro-CASP1 releasing CASP1 which further cleaves pro-IL1β into its mature form. The pro-apoptotic enzyme CASP10 and the adaptor CARD9 (acting downstream of C-type lectins signaling and upstream of the NF-κB pathway) are not represented in the diagram but were assayed for transcriptional regulation. Gene products, depicted with frames and colors were selected for microfluidic PCR assays, on basis of bibliographical data showing that they can be transcriptionally regulated following infection by RNA viruses. Furthermore, the selected genes sample most of the known signaling pathways triggered during a viral infectious process. P, phosphorylation.

**S1 Table. MERS-CoV N protein distribution in alpaca respiratory tracts by immunohistochemistry.**

**S2 Table. Sequence and characteristics of primers used for microfluidic qPCR assays in alpacas.**

**S3 Table. The expression levels of innate immune response genes, and viral loads (UpE and M mRNA) in MFPE nasal, tracheal and lung tissues.**

## References

1. Zaki AM, van Boheemen S, Bestebroer TM, Osterhaus ADME, Fouchier RAM. Isolation of a Novel Coronavirus from a Man with Pneumonia in Saudi Arabia. N Engl J Med. 2012;367: 1814–1820. doi:10.1056/nejmoa1211721

2. WHO | Middle East respiratory syndrome coronavirus (MERS-CoV). [cited 18 Jul 2020]. Available: https://www.who.int/emergencies/mers-cov/en/

3. Zumla A, Hui DS, Perlman S. Middle East respiratory syndrome. Lancet. 2015;386: 995–1007. doi:10.1016/S0140-6736(15)60454-8

4. Sabir JSM, Lam TT-Y, Ahmed MMM, Li L, Shen Y, E. M. Abo-Aba S, et al. Co-circulation of three camel coronavirus species and recombination of MERS-CoVs in Saudi Arabia. Science (80-). 2016;351: 81–84. doi:10.1126/science.aac8608

5. Adney DR, Bielefeldt-Ohmann H, Hartwig AE, Bowen RA. Infection, replication, and transmission of Middle East respiratory syndrome coronavirus in alpacas. Emerging Infectious Diseases. 2016. pp. 1031–1037. doi:10.3201/eid2206.160192

6. Crameri G, Durr PA, Klein R, Foord A, Yu M, Riddell S, et al. Experimental Infection and Response to Rechallenge of Alpacas with Middle East Respiratory Syndrome Coronavirus. Emerg Infect Dis. 2016;22: 1071–1074. doi:10.3201/eid2206.160007

7. Reusken CBEM, Schilp C, Raj VS, De Bruin E, Kohl RHG, Farag EABA, et al. MERS-CoV Infection of Alpaca in a Region Where MERS-CoV is Endemic. Emerging Infectious Diseases. 2016. pp. 1129–1131. doi:10.3201/eid2206.152113

8. David D, Rotenberg D, Khinich E, Erster O, Bardenstein S, van Straten M, et al. Middle East respiratory syndrome coronavirus specific antibodies in naturally exposed Israeli llamas, alpacas and camels. One Health. 2018. pp. 65–68. doi:10.1016/j.onehlt.2018.05.002

9. Vergara-Alert J, van den Brand JMA, Widagdo W, Muñoz M, Raj VS, Schipper D, et al. Livestock susceptibility to infection with middle east respiratory syndrome coronavirus. Emerging Infectious Diseases. 2017. pp. 232–240. doi:10.3201/eid2302.161239

10. Adney DR, Letko M, Ragan IK, Scott D, van Doremalen N, Bowen RA, et al. Bactrian camels shed large quantities of Middle East respiratory syndrome coronavirus (MERS-CoV) after experimental infection. Emerg Microbes Infect. 2019;8: 717–723. doi:10.1080/22221751.2019.1618687

11. Adney DR, van Doremalen N, Brown VR, Bushmaker T, Scott D, de Wit E, et al. Replication and shedding of MERS-CoV in upper respiratory tract of inoculated droamedary camels. Emerging Infectious Diseases. 2014. pp. 1999–2005. doi:10.3201/eid2012.141280

12. Reusken CBEM, Haagmans BL, Müller MA, Gutierrez C, Godeke G-J, Meyer B, et al. Middle East respiratory syndrome coronavirus neutralising serum antibodies in dromedary camels: a comparative serological study. Lancet Infect Dis. 2013;13: 859–866. doi:10.1016/S1473-3099(13)70164-6

13. Akira S, Takeda K, Kaisho T. Toll-like receptors: critical proteins linking innate and acquired immunity. Nat Immunol. 2001;2: 675–680. doi:10.1038/90609

14. Hartl D, Tirouvanziam R, Laval J, Greene CM, Habiel D, Sharma L, et al. Innate Immunity of the Lung: From Basic Mechanisms to Translational Medicine. J Innate Immun. 2018;10: 487–501. doi:10.1159/000487057

15. Alosaimi B, Hamed ME, Naeem A, Alsharef AA, AlQahtani SY, AlDosari KM, et al. MERS-CoV infection is associated with downregulation of genes encoding Th1 and Th2 cytokines/chemokines and elevated inflammatory innate immune response in the lower respiratory tract. Cytokine. 2020;126: 154895. doi:10.1016/j.cyto.2019.154895

16. Zhou J, Chu H, Li C, Wong BHY, Cheng ZS, Poon VKM, et al. Active replication of middle east respiratory syndrome coronavirus and aberrant induction of inflammatory cytokines and chemokines in human macrophages: Implications for pathogenesis. J Infect Dis. 2014;209: 1331–1342. doi:10.1093/infdis/jit504

17. Zhao X, Chu H, Wong BHY, Chiu MC, Wang D, Li C, et al. Activation of C-Type Lectin Receptor and (RIG)-I-Like Receptors Contributes to Proinflammatory Response in Middle East Respiratory Syndrome Coronavirus-Infected Macrophages. J Infect Dis. 2020;221: 647–659. doi:10.1093/infdis/jiz483

18. Kindler E, Jónsdóttir HR, Muth D, Hamming OJ, Hartmann R, Rodriguez R, et al. Efficient Replication of the Novel Human Betacoronavirus EMC on Primary Human Epithelium Highlights Its Zoonotic Potential. Buchmeier MJ, editor. MBio. 2013;4. doi:10.1128/mBio.00611-12

19. Zielecki F, Weber M, Eickmann M, Spiegelberg L, Zaki AM, Matrosovich M, et al. Human Cell Tropism and Innate Immune System Interactions of Human Respiratory Coronavirus EMC Compared to Those of Severe Acute Respiratory Syndrome Coronavirus. Journal of Virology. 2013. pp. 5300–5304. doi:10.1128/jvi.03496-12

20. Comar CE, Goldstein SA, Li Y, Yount B, Baric RS, Weiss SR. Antagonism of dsRNA-Induced Innate Immune Pathways by NS4a and NS4b Accessory Proteins during MERS Coronavirus Infection. Palese P, editor. MBio. 2019;10. doi:10.1128/mBio.00319-19

21. Chan RWY, Chu DKW, Poon LLM, Tao KP, Ng HY, Chan MCW, et al. Tropism and replication of Middle East respiratory syndrome coronavirus from dromedary camels in the human respiratory tract: An in-vitro and ex-vivo study. Lancet Respir Med. 2014;2: 813–822. doi:10.1016/S2213-2600(14)70158-4

22. Chu DKW, Hui KPY, Perera RAPM, Miguel E, Niemeyer D, Zhao J, et al. MERS coronaviruses from camels in Africa exhibit region-dependent genetic diversity. Proc Natl Acad Sci. 2018;115: 3144–3149. doi:10.1073/pnas.1718769115

23. Chan RWY, Chan MCW, Agnihothram S, Chan LLY, Kuok DIT, Fong JHM, et al. Tropism of and Innate Immune Responses to the Novel Human Betacoronavirus Lineage C Virus in Human Ex Vivo Respiratory Organ Cultures. Journal of Virology. 2013. pp. 6604–6614. doi:10.1128/jvi.00009-13

24. Niemeyer D, Zillinger T, Muth D, Zielecki F, Horvath G, Suliman T, et al. Middle East Respiratory Syndrome Coronavirus Accessory Protein 4a Is a Type I Interferon Antagonist. J Virol. 2013;87: 12489–12495. doi:10.1128/jvi.01845-13

25. Yang Y, Zhang L, Geng H, Deng Y, Huang B, Guo Y, et al. The structural and accessory proteins M, ORF 4a, ORF 4b, and ORF 5 of Middle East respiratory syndrome coronavirus (MERS-CoV) are potent interferon antagonists. Protein Cell. 2013;4: 951–961. doi:10.1007/s13238-013-3096-8

26. Menachery VD, Schäfer A, Sims AC, Gralinski LE, Long C, Webb-Robertson B-J, et al. Pathogenic influenza viruses and coronaviruses utilize similar and contrasting approaches to control interferon-stimulated gene responses. MBio. 2014;5: 1–11. doi:10.1128/mBio.01174-14

27. Lau SKP, Lau CCY, Chan KH, Li CPY, Chen H, Jin DY, et al. Delayed induction of proinflammatory cytokines and suppression of innate antiviral response by the novel Middle East respiratory syndrome coronavirus: Implications for pathogenesis and treatment. Journal of General Virology. 2013. pp. 2679–2690. doi:10.1099/vir.0.055533-0

28. Matthews KL, Coleman CM, van der Meer Y, Snijder EJ, Frieman MB. The ORF4b-encoded accessory proteins of Middle East respiratory syndrome coronavirus and two related bat coronaviruses localize to the nucleus and inhibit innate immune signalling. J Gen Virol. 2014;95: 874–882. doi:10.1099/vir.0.062059-0

29. Rabouw HH, Langereis MA, Knaap RCM, Dalebout TJ, Canton J, Sola I, et al. Middle East Respiratory Coronavirus Accessory Protein 4a Inhibits PKR-Mediated Antiviral Stress Responses. Frieman MB, editor. PLOS Pathog. 2016;12: e1005982. doi:10.1371/journal.ppat.1005982

30. Chang C-Y, Liu HM, Chang M-F, Chang SC. Middle East Respiratory Syndrome Coronavirus Nucleocapsid Protein Suppresses Type I and Type III Interferon Induction by Targeting RIG-I Signaling. Gallagher T, editor. J Virol. 2020;94. doi:10.1128/JVI.00099-20

31. Lazear HM, Schoggins JW, Diamond MS. Shared and Distinct Functions of Type I and Type III Interferons. Immunity. Cell Press; 2019. pp. 907–923. doi:10.1016/j.immuni.2019.03.025

32. Ahn M, Anderson DE, Zhang Q, Tan CW, Lim BL, Luko K, et al. Dampened NLRP3-mediated inflammation in bats and implications for a special viral reservoir host. Nature Microbiology. 2019. pp. 789–799. doi:10.1038/s41564-019-0371-3

33. El-Kafrawy SA, Corman VM, Tolah AM, Al Masaudi SB, Hassan AM, Müller MA, et al. Enzootic patterns of Middle East respiratory syndrome coronavirus in imported African and local Arabian dromedary camels: a prospective genomic study. Lancet Planet Heal. 2019;3: e521–e528. doi:10.1016/S2542-5196(19)30243-8

34. Widagdo W, Raj VS, Schipper D, Kolijn K, van Leenders GJLH, Bosch BJ, et al. Differential Expression of the Middle East Respiratory Syndrome Coronavirus Receptor in the Upper Respiratory Tracts of Humans and Dromedary Camels. J Virol. 2016;90: 4838–4842. doi:10.1128/JVI.02994-15

35. Channappanavar R, Fehr AR, Zheng J, Wohlford-Lenane C, Abrahante JE, Mack M, et al. IFN-I response timing relative to virus replication determines MERS coronavirus infection outcomes. J Clin Invest. 2019;129: 3625–3639. doi:10.1172/JCI126363

36. Klinkhammer J, Schnepf D, Ye L, Schwaderlapp M, Gad HH, Hartmann R, et al. IFN-λ prevents influenza virus spread from the upper airways to the lungs and limits virus transmission. Elife. 2018;7. doi:10.7554/eLife.33354

37. Shokri S, Mahmoudvand S, Taherkhani R, Farshadpour F. Modulation of the immune response by Middle East respiratory syndrome coronavirus. Journal of Cellular Physiology. Wiley-Liss Inc.; 2019. pp. 2143–2151. doi:10.1002/jcp.27155

38. Alsaad KO, Hajeer AH, Al Balwi M, Al Moaiqel M, Al Oudah N, Al Ajlan A, et al. Histopathology of Middle East respiratory syndrome coronovirus (MERS-CoV) infection – clinicopathological and ultrastructural study. Histopathology. 2018;72: 516–524. doi:10.1111/his.13379

39. Ouyang W, Rutz S, Crellin NK, Valdez PA, Hymowitz SG. Regulation and Functions of the IL-10 Family of Cytokines in Inflammation and Disease. Annu Rev Immunol. 2011;29: 71–109. doi:10.1146/annurev-immunol-031210-101312

40. Hemann EA, Green R, Turnbull JB, Langlois RA, Savan R, Gale M. Interferon-λ modulates dendritic cells to facilitate T cell immunity during infection with influenza A virus. Nat Immunol. 2019;20: 1035–1045. doi:10.1038/s41590-019-0408-z

41. Scheuplein VA, Seifried J, Malczyk AH, Miller L, Höcker L, Vergara-Alert J, et al. High Secretion of Interferons by Human Plasmacytoid Dendritic Cells upon Recognition of Middle East Respiratory Syndrome Coronavirus. J Virol. 2015;89: 3859–3869. doi:10.1128/jvi.03607-14

42. Mascarell L, Lombardi V, Louise A, Saint-Lu N, Chabre H, Moussu H, et al. Oral dendritic cells mediate antigen-specific tolerance by stimulating TH1 and regulatory CD4+ T cells. J Allergy Clin Immunol. 2008;122: 603–609.e5. doi:10.1016/j.jaci.2008.06.034

43. Ali S, Mann-Nüttel R, Schulze A, Richter L, Alferink J, Scheu S. Sources of Type I Interferons in Infectious Immunity: Plasmacytoid Dendritic Cells Not Always in the Driver’s Seat. Front Immunol. 2019;10. doi:10.3389/fimmu.2019.00778

44. Krausgruber T, Blazek K, Smallie T, Alzabin S, Lockstone H, Sahgal N, et al. IRF5 promotes inflammatory macrophage polarization and TH1-TH17 responses. Nat Immunol. 2011;12: 231–238. doi:10.1038/ni.1990

45. Weiss M, Blazek K, Byrne AJ, Perocheau DP, Udalova IA. IRF5 Is a Specific Marker of Inflammatory Macrophages In Vivo. Mediators Inflamm. 2013;2013: 1–9. doi:10.1155/2013/245804

46. Verbist KC, Klonowski KD. Functions of IL-15 in anti-viral immunity: Multiplicity and variety. Cytokine. 2012;59: 467–478. doi:10.1016/j.cyto.2012.05.020

47. Guarda G, Zenger M, Yazdi AS, Schroder K, Ferrero I, Menu P, et al. Differential Expression of NLRP3 among Hematopoietic Cells. J Immunol. 2011;186: 2529–2534. doi:10.4049/jimmunol.1002720

48. Shao B-Z, Xu Z-Q, Han B-Z, Su D-F, Liu C. NLRP3 inflammasome and its inhibitors: a review. Front Pharmacol. 2015;6. doi:10.3389/fphar.2015.00262

49. Hayden MS, Ghosh S. Signaling to NF-κB. Genes and Development. Cold Spring Harbor Laboratory Press; 2004. pp. 2195–2224. doi:10.1101/gad.1228704

50. Chan JFW, Yao Y, Yeung ML, Deng W, Bao L, Jia L, et al. Treatment with lopinavir/ritonavir or interferon-β1b improves outcome of MERSCoV infection in a nonhuman primate model of common marmoset. J Infect Dis. 2015;212: 1904–1913. doi:10.1093/infdis/jiv392

51. Arabi YM, Shalhoub S, Mandourah Y, Al-Hameed F, Al-Omari A, Al Qasim E, et al. Ribavirin and Interferon Therapy for Critically Ill Patients With Middle East Respiratory Syndrome: A Multicenter Observational Study. Clin Infect Dis. 2020;70: 1837–1844. doi:10.1093/cid/ciz544

